# A Simplest Bioinformatics Pipeline for Whole Transcriptome Sequencing: Overview of the Processing and Steps from Raw Data to Downstream Analysis

**DOI:** 10.1101/836973

**Authors:** Ayam Gupta, Sonal Gupta, Suresh Kumar Jatawa, Ashwani Kumar, Prashanth Suravajhala

**Affiliations:** Department of Biotechnology and Bioinformatics, Birla Institute of Scientific Research, Jaipur 302001, Rajasthan, India; Department of Pediatrics, SMS Medical College and Hospital, Jaipur, Rajasthan, India; School of Biotechnology, Rajiv Gandhi Proudyogiki Vishwavidyalaya, State Technological University of Madhya Pradesh, Bhopal, Madhya Pradesh, India; DNA Xperts, New Delhi, India; Bioclues.org, India

**Keywords:** RNA-Seq, Transcriptome Sequencing, Bioinformatics pipeline, reinstantiation

## Abstract

Recent advances in next generation sequencing (NGS) technologies have heralded the genomic research. From the good-old inferring differentially expressed genes (DEG) using microarray to the current adage NGS-based whole transcriptome or RNA-Seq pipelines, there have been advances and improvements. With several bioinformatics pipelines for analysing RNA-Seq on rise, inferring the candidate DEGs prove to be a cumbersome approach as one may have to reach consensus among all the pipelines. To Check this, we have benchmarked the well known cufflinks-cuffdiff pipeline on a set of datasets and outline it in the form of a protocol where researchers interested in performing whole transcriptome shotgun sequencing and it’s downstream analysis can better disseminate the analysis using their datasets.

## Background

Whole transcriptome shotgun sequencing (WTSS or RNA-seq) allows us to discover new genes and transcripts and measure their expression patterns in a single assay (Trapnell *et al.*, 2012). The quantitative analysis of gene expression allows the researchers in predicting the molecular mechanisms underlying genome regulation. In addition, RNA-seq can also detect lowly expressed transcripts in comparison to microarray based expression quantification (Illumina, 2011; Nookaew *et al.*, 2012; Zhao *et al.*, 2014). While it enables us to characterize RNA transcripts in various organisms including both the coding and non-coding part, it has helped researchers in identifying differential expression patterns in cases Vs. control samples. The RNA-seq analyses is analyzed with robust and sensitive algorithms; even as RNA-seq pipeline, as a protocol would vary based on how good the publicly available datasets were used for an efficient benchmarking. In this protocol, we present a bioinformatics workflow for quantitative analysis of RNA-seq data using cufflinks-cuffmerge-cuffdiff which includes open source tools all through the quality check to ascertaining differentially expressed genes (see Software section).

### Equipment

#### Computer

Ideal configuration: 33/64GB RAM with 64 core CPUs in an Ubuntu operating system (16.04.4 LTS machine)

### Software

All the softwares were downloaded/used from the following locations:

1. FastQC- https://www.bioinformatics.babraham.ac.uk/projects/fastqc
2. Bowtie2- https://bowtie-bio.sourceforge.net/bowtie2
3. HISAT2- https://github.com/infphilo/hisat2
4. Samtools- http://samtools.sourceforge.net
5. Cufflinks- https://github.com/cole-trapnell-lab/cufflinks
6. DeSeq2- https://bioconductor.org/packages/release/bioc/html/DESeq2.html
7. Feature Counts- apt-get install feature counts
8. 1000 genomes dataset- http://www.internationalgenome.org/

### Datasets

For benchmarking, we have downloaded publicly available datasets from 1000 Genomes datasets (http://www.internationalgenome.org/) based on three different populations *viz*. East Asian (ERR1050076 and ERR1050078), American (HG00731 and HG00733) and African ancestory (NA19238 and NA19240) (Table 1). Each population contains individual as parent and child and type of RNA sequencing is strand specific using Illumina platform. The files were downloaded either using FTP or fasterq-dump (https://www.internationalgenome.org/data-portal/sample/) (Durbin *et al.*, 2010)

**Table 1:**
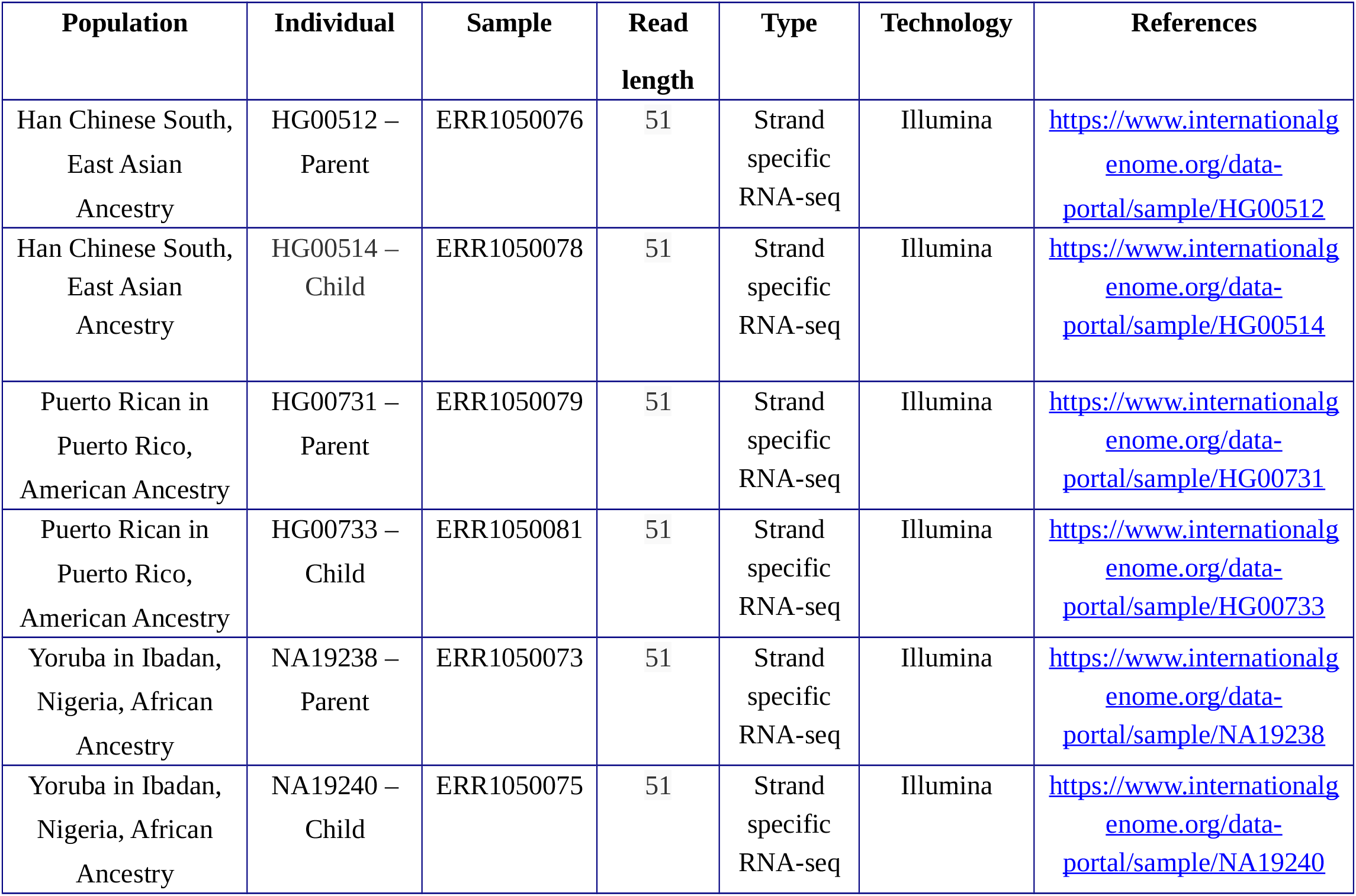
1000 genomes samples used for benchmarking

### Procedure

The raw file (fastq) was processed using various steps which included quality check, alignment, sorting and annotation. The pipeline is an amalgamation of various tools, *viz*. FastQC, Bowtie2/HISAT2, samtools, cufflinks along with imperative files containing the human genome reference sequence (hg38 from UCSC human genome browser) and genes coordinate information file. Since the commands in pipeline run on linux, all commands are presumptively case sensitive. This pipeline was run on a 64 GB RAM with 64 core CPUs in an Ubuntu operating system (16.04.4 LTS machine), but this can also be run on a minimum 16 GB RAM machine depending upon the raw fastq file size. A shell script was created with extension .sh and all the commands with details are mentioned below.

#### A. Preprocessing of raw data

##### Quality check

The data analysis of NGS studies rely on the raw data as it delivers a prompt insight of the sequence quality for further downstream analysis. In our pipeline, we used FastQC as it plots the read depth and quality score besides a host of other statistical inferences.

1. ./fastqc ~/samples/sample1.fastq

**Figure 1.**
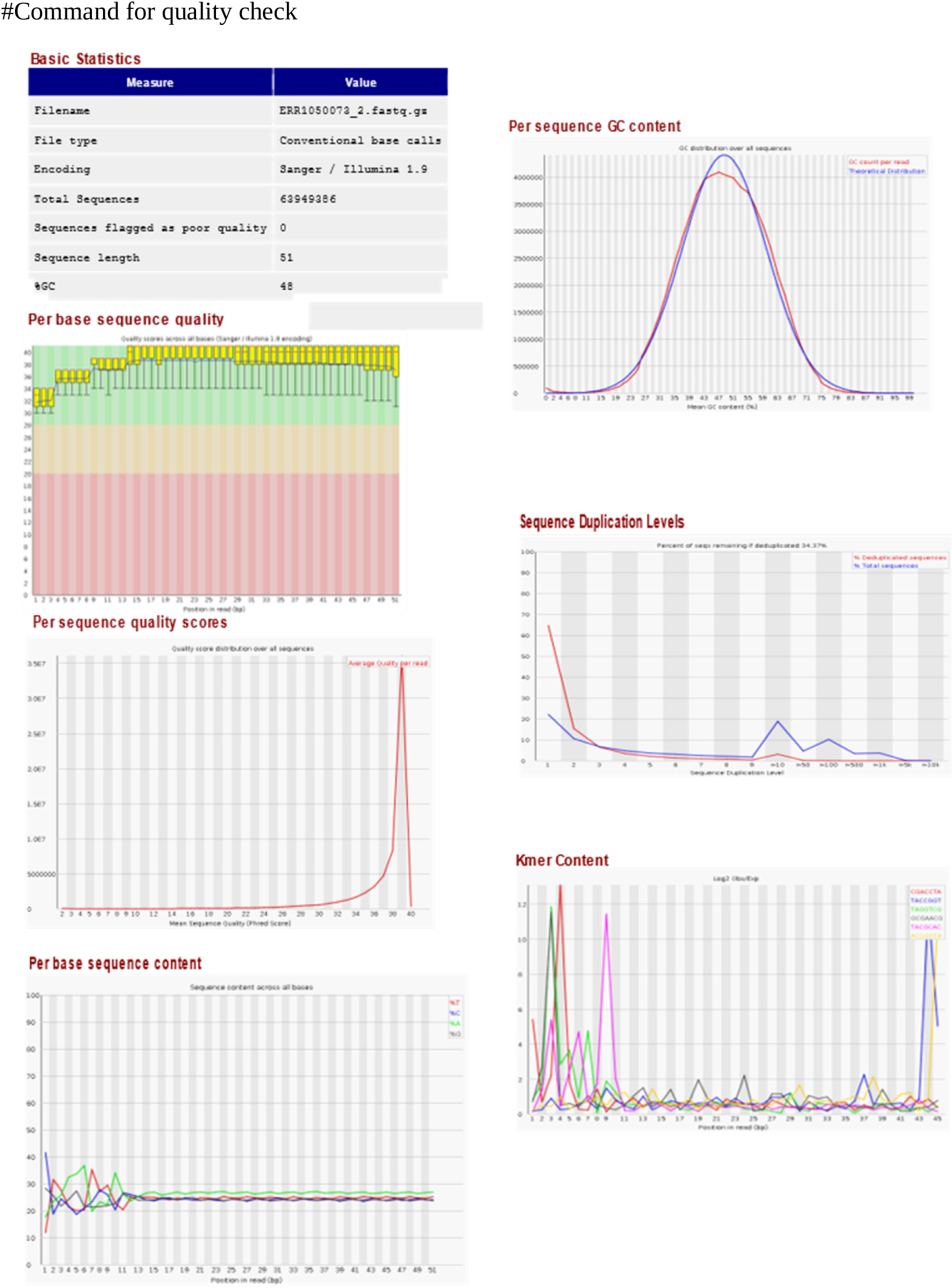
Output of the box plots and figures generated by FastQC run, containing information on statistics, quality, read coverage, depth, yield, based per read call, *etc*.

###### Indexing human genome

Bowtie2 and HISAT2 were used to index reference genome both of which work at high speed and are memory efficient.When compared to Bowtie2, the HISAT2 indexing is more memory efficient although the latter cannot compare the intron splice junctions.

./bowtie2-build reference_in bt2_index_base

./hisat2-build reference_in ht2_index_base

#builds indices which are required for aligning reads

When the aforementioned HISAT2 commands are run, hisat2-build outputs a set of 8 files (with suffixes .1.ht2, .2.ht2, .3.ht2, .4.ht2, .5.ht2, .6.ht2, .7.ht2, and .8.ht2). These files together constitute the index that is needed to align reads to the reference genome. The original sequence FASTA files are no longer used by HISAT2 once the index is built. In case of Bowtie2, the current directory will contain six new files that all start with reference and end with .1.bt2, .2.bt2, .3.bt2, .4.bt2, .rev.1.bt2, and .rev.2.bt2. While the first four files are the forward strands, the rev files indicate the reverse indexed sequences in case of bowtie2.

##### Alignment and post processing

HISAT2 is fast and sensitive alignment used to map RNA-seq reads to the genome. Like Bowtie, HISAT uses burrows-wheeler transform (BWT) to compress the genomes so that they require very little memory to store. Unlike BWA and Bowtie, HISAT2 builds a whole genome global index and thousands of small local indexes to make spliced alignment possible to run on a standard system in an efficient way (Kim *et al.*, 2015). Bowtie2 is used for short read alignment and uses very little RAM with accuracy and modest performance in indexing the alignment (Langmead and Salzberg, 2012). The mismatch or any sequencing errors or small genetic variation between samples and reference genome could be checked using the following commands:

3. ./hisat2 –x reference_hisat_filename −1 path/filename1 −2 path/filename2 > filename.sam

./bowtie2 -x reference_filename −1 path/filename1 −2 path/filename2 > filename.sam

*Note: The -2 option may be omitted for single-end sequences and –U may be used instead.

HISAT2 and Bowtie2 align a set of unpaired reads (in fastq or .fq format) to the reference genome using the Ferragina and Manzini (FM)-index (Langmead and Salzberg, 2012). The alignment results output in SAM format (Li *et al.*, 2009) and a short alignment summary is written to the console.

**Samtools** is a collection of tools to manipulate the resulting alignment in SAM/BAM format. SAM stands for sequence alignment/map format and its corresponding format is binary mapped format (BAM). SAM could be converted into different alignment format, sort, merge, alignment, remove duplicates, call SNPs and short indel variants; whereas the BAM and following indices (.bai) are used to view the index of the binary aligned sequences. The basic necessity of having the binary mapped files is to save the memory.

4. ./samtools import reference.fa ~/samples/sample1.sam > sample1.bam

**Sorting BAM**: A sorted BAM file is used for streamlined data processing to avoid extra alignment when loaded into memory. Whereas it can be easily indexed for faster retrieval of alignment, overlapping is needed to retrieve alignment from a region. All BAM files needs to be sorted and indexed for better performance. Samtools sort command is used to convert the BAM file to a sorted BAM file and samtools index to index BAM file.

5. ~/samtools sort ~/samples/sample1.bam -o sample1.sorted

6. ~/samtools index ~/samples/sample1.sorted

#### B. Examining differentially expressed genes

##### Cufflinks

The Cufflinks suite of tools is used to perform a number of analyses for RNA-Seq experiments. It assembles the transcripts and estimates their abundances, and finally examines DEGs from the samples. The Cufflinks suite includes different programs that work together to perform these analyses.

1. ./cufflinks in.sorted.bam -o sample.out

It is included in Cufflinks package which helps in merging the assemblies. The merged assemblies provides us a basis for calculating genes and transcripts expression in each condition which are finally analysed for DEGs through Cuffdiff command. Cuffmerge, on the other hand is preferentially used if there are replicate samples.

2. ./cuffmerge -g reference.gtf -s reference.fasample.merged

**Cuffdiff** is a part of the Cufflinks suite. It takes and compares the aligned reads from RNA-seq samples from two or more conditions and identifies transcripts that are differentially expressed using a rigorous statistical analysis, *viz*. from the log2fold change which are then reported as set of text files and can be plotted graphically of our choice.

3. ./cuffdiff ~/samples/sample1.out/transcripts.gtf ~/samples/sample1.sorted ~/samples/reference_ genome >sample_result

#### C. Exploring differential expression with DeSeq2 package

**Feature counts** is a program for assigning mapping reads to genomic features such as genes, exons, promoters and genomic bins (Yang *et al.*, 2014)..

1. featureCounts -B -p -g gene_id -a ~/reference.gtf -o ~/sample.sorted.bam ~/sample.sorted.bam

**DESeq2,** a Bioconductor package available in R is used for DEG analysis based on the negative binomial distribution.

1. **Alignment of Sequencing Reads** After benchmarking, we got a good recovery rate for DEGs across these tools. From six samples, we obtained a RNA-seq quality checked using FastQC. The reads were aligned to the human reference genome assembly (hg38) and consensus was reached using Bowtie2 and HISAT2. While the average alignment percentage of 91.82 was observed for Bowtie2, for HISAT2, it was found to be 97.27% even as the percentage of reads mapped to the reference genome was found to be similar between all six groups suggesting that there were no sequencing biases in the data (Table 2). Further from coverage percentage analysis between DESeq2 pipeline and our pipeline we observed that Bowtie2 and HISAT2 generated feature counts from DESeq2, ca. 60% and 50% coverage respectively. This is much lower than Bowtie2 (~90%) and HISAT2 (>90%) although it generated alignment files from our pipeline across all the samples (Figure 2).
2. **DEG analyses** The files were processed using cufflinks software with alternating the transcripts.out file from both the samples of the same population (Figure 3). Cuffdiff analysis was done using transcripts.out file (output file from cuffdiff) with parent:child as well as child:parent which yielded different results. We obtained an average of 870 (YRI), 928 (PUR) and 800 (CHS) DEGs using cuffdiff taken as parent: child and while taken as child: parent we obtained an average of 3036 (YRI), 2963 (PUR) and 3093 (CHS) DEGs.

**Table 2:** Alignment coverage obtained using Bowtie2 and Hisat2 aligners.

| SAMPLE | BOWTIE2 | HISAT2 |
| --- | --- | --- |
| ERR1050076 | 91.91% | 97.49% |
| ERR1050078 | 92.07% | 97.05% |
| ERR1050079 | 91.87% | 97.29% |
| ERR1050081 | 91.64% | 97.27% |
| ERR1050073 | 91.66% | 97.45% |
| ERR1050075 | 91.75% | 97.09% |

**Figure 2:**
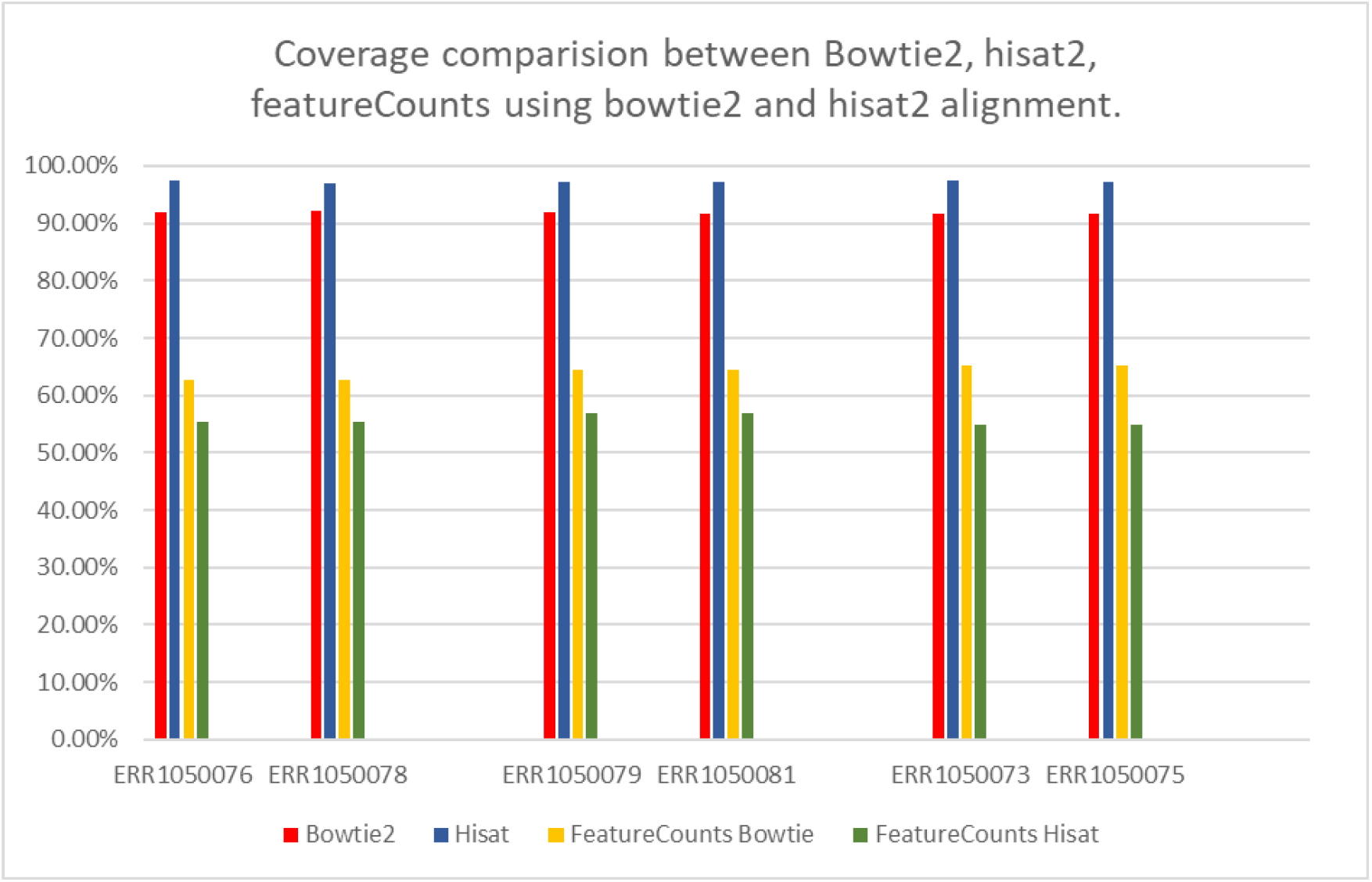
Coverage comparison between Bowtie2, Hisat2 and Subread packages.

**Figure 3:**
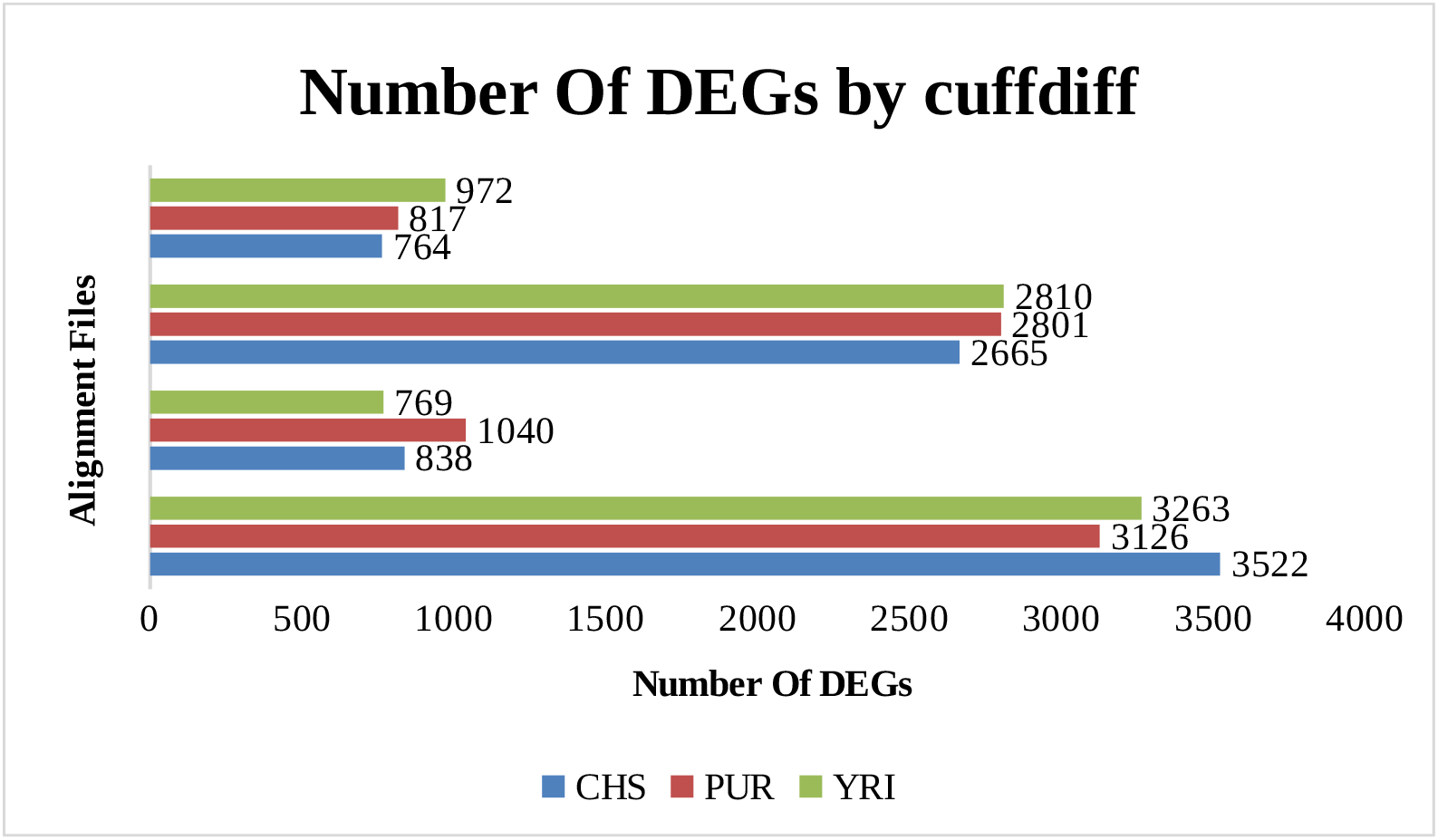
Number of DEGs with cufflinks tool uses alignment files from Hisat2 and Bowtie2.

**Figure 4:**
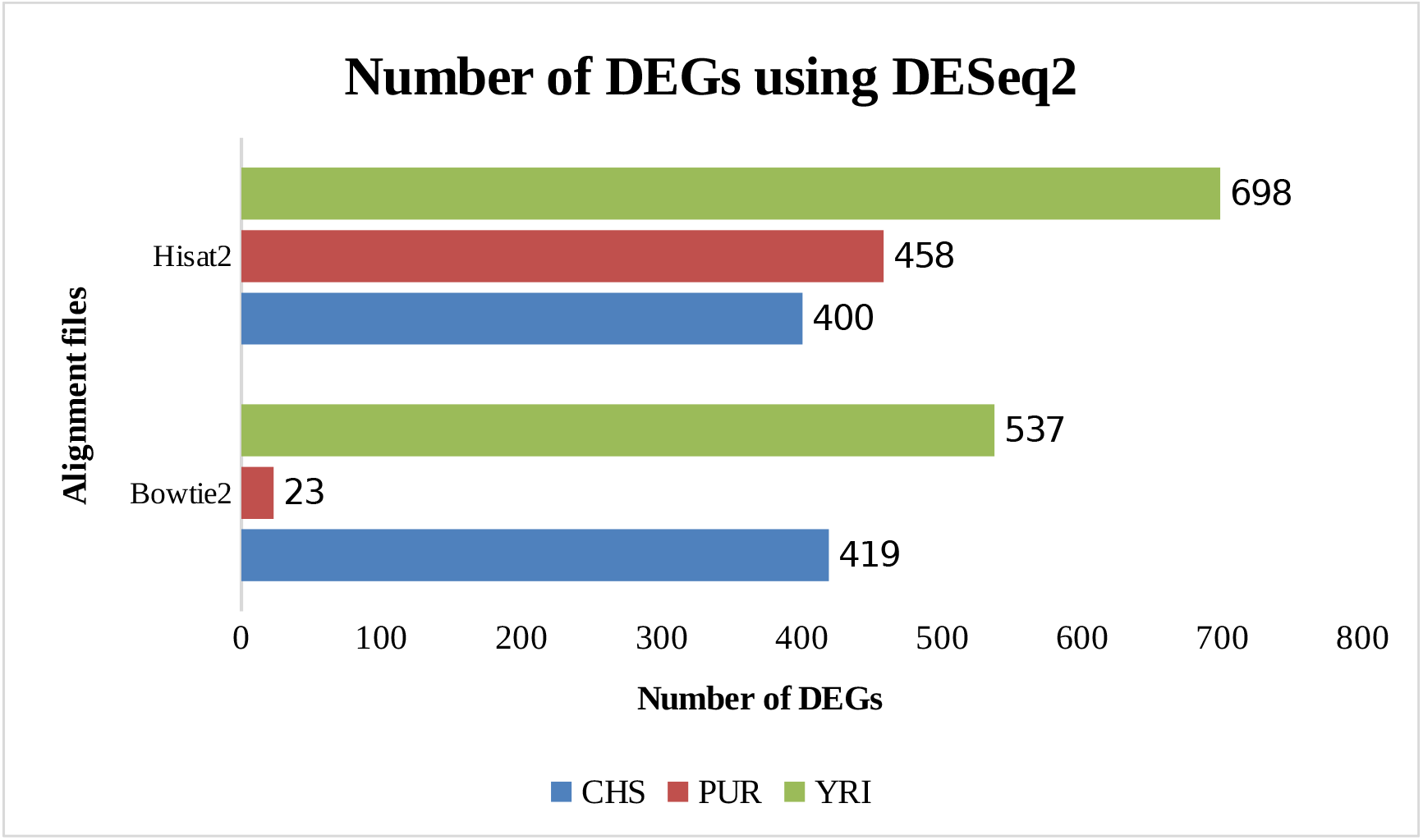
Total number of DEGs discovered using the DESeq2 pipeline.

**Figure 5:**
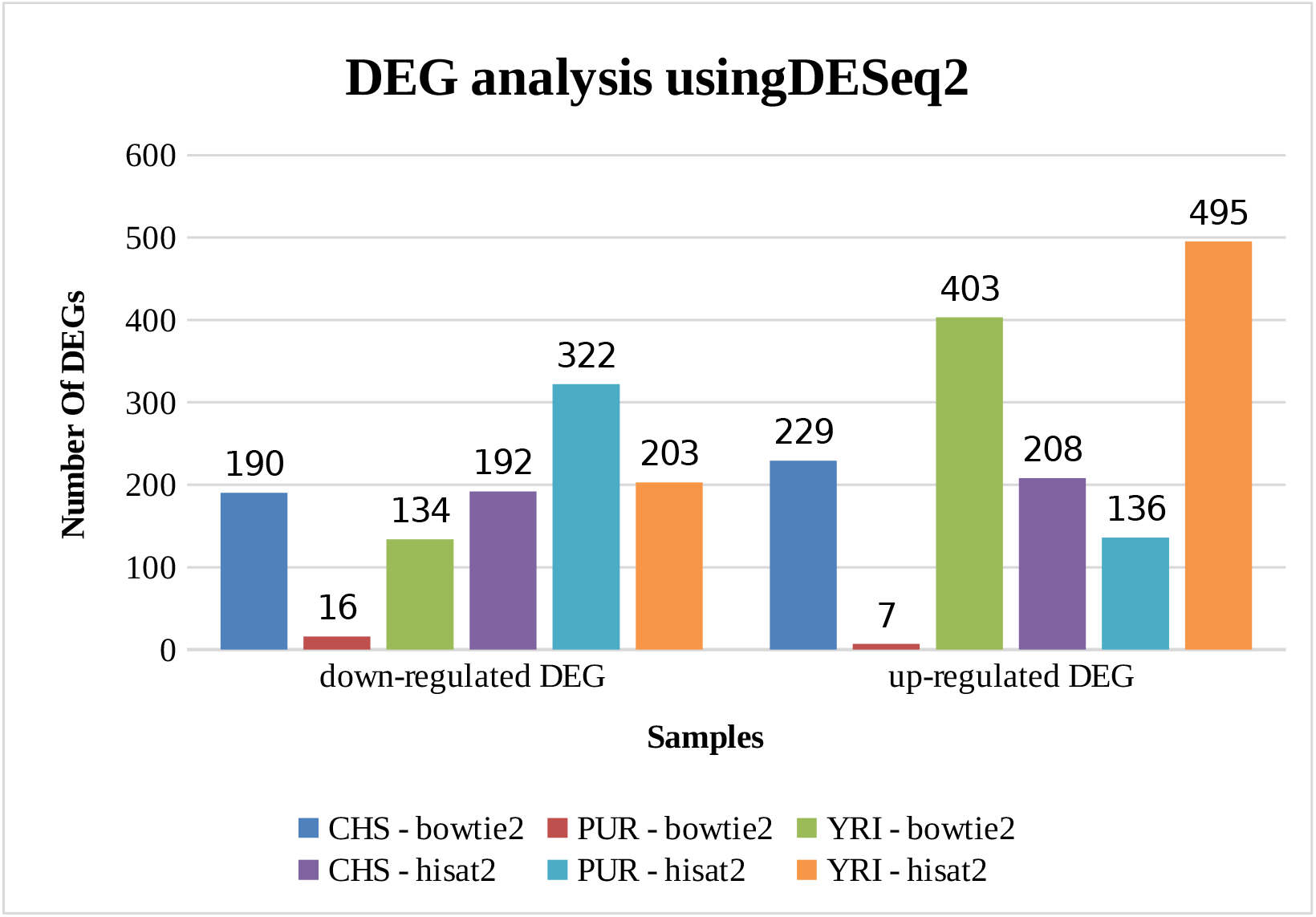
Down regulated and up-regulated gene count using the DESeq2 pipeline. *No normalization is done in either of the analysis.

**Figure 6:**
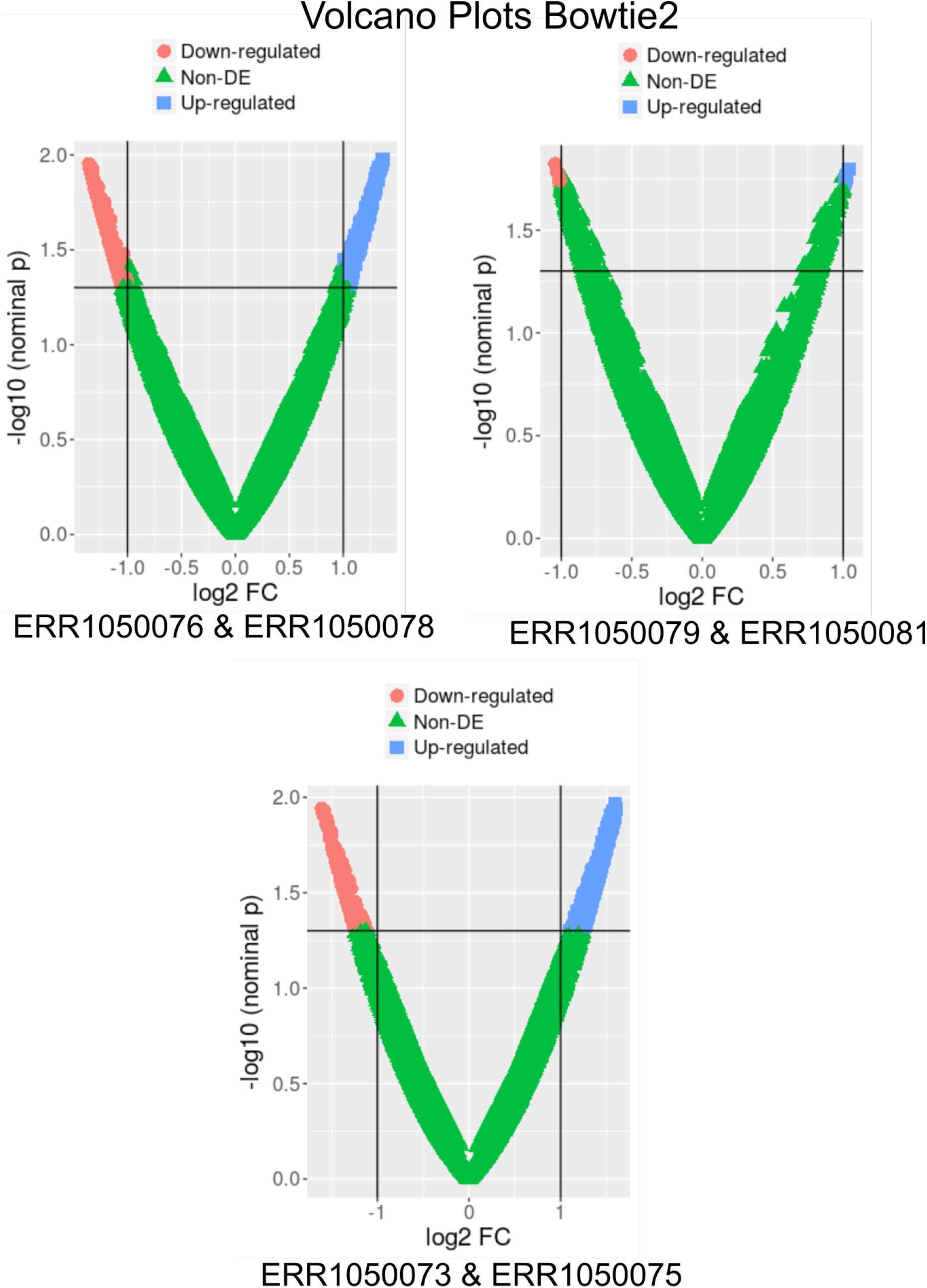
Volcano plots from DESeq2 analysis showing log2fold (DEGs) change in the samples aligned using Bowtie2

**Figure 7:**
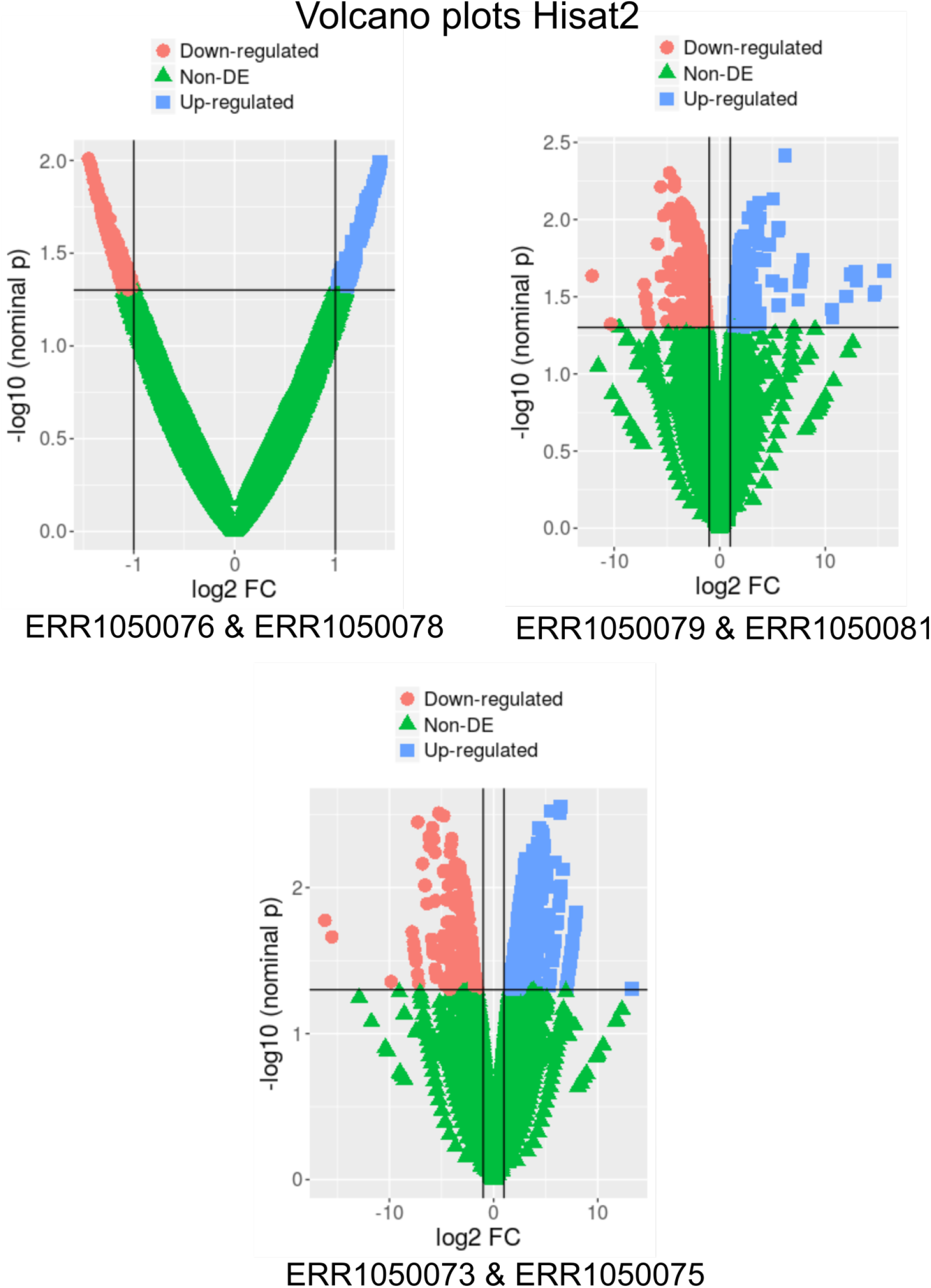
Volcano plots from DESeq2 analysis showing log2fold (DEGs) change in the samples aligned using Hisat2.

## Conclusions

Benchmarking is a gold standard for any NGS pipeline. It gives us inherent statistical interpretation of how the data is used for further analyses. In this protocol, we have presented a RNA-Seq pipeline to screen the DEGs and we hope that this pipeline could be aptly used by beginners. While the cufflinks-cuffdiff pipeline is a simplest approach to identify the DEGs, the tools such as DESeq serves as another alternative.

